# Genetic landscape of recessive diseases in the Vietnamese population from large-scale clinical exome sequencing

**DOI:** 10.1101/2020.10.27.358358

**Authors:** Ngoc Hieu Tran, Thanh-Huong Nguyen Thi, Hung-Sang Tang, Le-Phuc Hoang, Trung-Hieu Le Nguyen, Nhat-Thang Tran, Thu-Huong Nhat Trinh, Van Thong Nguyen, Bao-Han Huu Nguyen, Hieu Trong Nguyen, Loc Phuoc Doan, Ngoc-Minh Phan, Kim-Huong Thi Nguyen, Hong-Dang Luu Nguyen, Minh-Tam Thi Quach, Thanh-Phuong Thi Nguyen, Vu Uyen Tran, Dinh-Vinh Tran, Quynh-Tho Thi Nguyen, Thanh-Thuy Thi Do, Nien Vinh Lam, Phuong Cao Thi Ngoc, Dinh Kiet Truong, Hoai-Nghia Nguyen, Minh-Duy Phan, Hoa Giang

## Abstract

**Purpose:** Accurate profiling of population-specific recessive diseases is essential for the design of cost-effective carrier screening programs. However, minority populations and ethnic groups, including Vietnamese, are still under-represented in existing genetic studies. Here we reported the first comprehensive study of recessive diseases in the Vietnamese population.

**Methods:** Clinical exome sequencing (CES) data of 4,503 disease-associated genes obtained from a cohort of 985 Vietnamese individuals was analyzed to identify pathogenic variants, associated diseases and their carrier frequencies in the population.

**Results:** Eighty-five recessive diseases were identified in the Vietnamese population, among which seventeen diseases had carrier frequencies of at least 1% (1 in 100 individuals). Three diseases were especially prevalent in the Vietnamese population with carrier frequencies of 2-12 times higher than in other East Asia or the world populations, including Beta-thalassemia (1 in 25), citrin deficiency (1 in 33) and phenylketonuria (1 in 40). Seven novel pathogenic and three likely pathogenic variants associated with nine recessive diseases were also discovered.

**Conclusions:** The comprehensive profile of recessive diseases identified in this study shall enable the design of cost-effective carrier screening programs specific to the Vietnamese population. The newly discovered pathogenic variants may also exist in other populations at extremely low frequencies, thus representing a valuable resource for future research. Our study has demonstrated the advantage of population-specific genetic studies to advance the knowledge and practice of medical genetics.

## INTRODUCTION

As high-throughput sequencing technologies are getting more popular and affordable, carrier screening has become a routine, essential tool for preventive healthcare and offers a great resource of information to guide public health policies [1–3]. Individuals identified as carriers of pathogenic genes and associated disorders may take preventive steps to reduce the risk of having their offspring inherit these disorders, such as preimplantation genetic diagnosis of embryos and/or early prenatal genetic testing. In addition, newborn screening programs enable early diagnosis and effective treatment of affected children, which not only significantly improve the outcomes but also reduce the treatment costs and efforts.

However, there are more than a thousand of Mendelian inherited disorders that have been documented, according to the Online Mendelian Inheritance in Man (OMIM) database [4]. Most of the diseases are rare and their prevalence depends heavily on specific populations and ethnicities. Thus, accurate profiling of population- or ethnicity-specific inherited diseases is essential for the design of cost-effective and comprehensive carrier screening programs. For example, cystic fibrosis is recommended for carrier screening for individuals of Caucasian or Ashkenazi Jewish ancestry, Tay–Sachs disease for individuals of Ashkenazi Jewish ancestry, and Beta-thalassemia for individuals from Mediterranean regions [1]. The critical problem, however, is the under-representation of minority populations and ethnic groups in existing genetic studies and databases [5–7]. Gurdasani *et al.* found that nearly 78% of the participants in genetic studies had European ancestries, whereas the two major populations, Asian and African, only accounted for 11% and 2.4%, respectively [6]. Indeed, genetic research on the Vietnamese population still lags behind other Western and Asian populations, despite recent efforts of genome sequencing projects in the country [8, 9]. There has been no research to study the prevalence of inherited disorders in the Vietnamese population. Even for well-known diseases in the population such as non-syndromic hearing loss and deafness or Beta-thalassemia, their exact carrier frequencies and associated pathogenic variants are still unknown. Carrier screening in Vietnam is still in its infancy.

In this study, we reported the first comprehensive profile of 85 recessive diseases and their prevalence in the Vietnamese population. The study was performed on the clinical exome sequencing data of 4,503 genes obtained from a cohort of 985 Vietnamese individuals. We analyzed the genetic variants obtained from these individuals and identified all pathogenic variants, genes, associated diseases, and their carrier frequencies in the Vietnamese population. We also compared the results to other populations and highlighted three diseases that were found to be specific to the Vietnamese population. Finally, we identified seven novel pathogenic and three likely pathogenic variants and discussed how they might cause severe damages in nine associated diseases. Our study made an important step to advance the practice of medical genetics in Vietnam by providing the first inclusive picture of recessive diseases in the population and facilitating the development of carrier screening programs in the country.

## MATERIALS AND METHODS

### Recruitment of study participants

In this study, 985 individuals were recruited from 51 hospitals and clinics across Vietnam. The participants have approved and given written informed consent to the anonymous re-use of their genomic data for this study. The data was de-identified and aggregated for genetic analysis of the Vietnamese population. The study was approved by the institutional ethics committee of the University of Medicine and Pharmacy, Ho Chi Minh city, Vietnam

### Gene panel

Targeted exome sequencing with a panel of 4,503 clinically relevant genes was performed to study inherited diseases in the Vietnamese population. The full list of genes is provided in Supplementary Table S1.

### Clinical exome sequencing

Libraries were prepared from 2 ng of DNA using the NEBNext Ultra II FS DNA library prep kit (New England Biolabs, USA) following the manufacturer’s instructions. Subsequently, libraries were pooled prior to hybridization with the xGen Lockdown probes for 4,503 targeted genes (Integrated DNA Technologies, USA). Exome sequencing was performed using NextSeq 500/550 High output kits v2 (150 cycles) on Illumina NextSeq 550 system (Illumina, USA) with the coverage of 100x

### Variant calling and analysis

Quality control and alignment of sequencing data to the human reference genome (build GRCh38) was performed following an established analysis workflow with FastQC [10], trimmomatic [11], bwa [12], samtools [13], and bedtools [14]. Variant calling was performed using GATK 3.8, followed by standard filters of quality and sequencing coverage [15]. We also filtered out variants with allele frequencies less than 0.1% and variants that were located outside of the target regions of our gene panel. The final variant call set was annotated against dbSNP (version 151, [16]) and ClinVar (version 20191231, [17]) databases, and was analyzed for their potential consequences using VEP [18]. Principal component analysis was performed using PLINK (version 1.9, [19]).

## RESULTS

### Study cohort

The cohort in our study included 985 participants who were recruited from 51 hospitals and clinics across Vietnam. The ages and types of samples of the participants are summarized in Table 1. The average age was 23.8 weeks gestational for fetuses, 4.4 years for children (54% male, 46% female), and 39.5 years for adults (43% male, 57% female). Among types of samples, most were blood (65.7%), followed by amniotic fluid (18.3%), buccal swab (12.1%), placental (1.32%), umbilical cord (0.8%) and others (1.83%).

**Table 1.** Summary of participants (n=985)

| Types of ages | Types of samples |  |  |  |  |  | OVERALL TOTAL |
| --- | --- | --- | --- | --- | --- | --- | --- |
|  | Blood | Amniotic fluid | Placental | Umbilical cord | Buccal swab | Others |  |
| Fetus | 0 | 180 | 13 | 8 | 0 | 0 | 201 (20.4%) |
| Child (age<18) | 212 | 0 | 0 | 0 | 90 | 5 | 307 (31.2%) |
| Adult (age≥18) | 435 | 0 | 0 | 0 | 29 | 13 | 477 (48.4%) |
| OVERALL TOTAL | 647 (65.7%) | 180 (18.3%) | 13 (1.3%) | 8 (0.8%) | 119 (12.1%) | 18 (1.8%) |  |

### Summary of genetic variants in the study cohort

The aggregated variant call set from 985 individuals, denoted as G4500, consisted of 67,140 variants, including 61,327 SNPs (91.3%) and 5,813 indels (8.7%). Figures 1a-c show the comparison of the G4500 call set to that of the KHV population (Kinh in Ho Chi Minh City, Vietnam) from the 1000 genomes project [7] and the dbSNP database (for this comparison, we only considered variants located within the target regions of our gene panel). We found that 27,655 variants (41.2%) of the G4500 call set had been reported earlier in the KHV call set (Figure 1a), and their allele frequencies were consistent between the two call sets with a strong Pearson correlation of 99.0% (Figure 1c). We also noted that the G4500 call set missed 8,634 KHV variants, and further investigation showed that most (91.7%) of those variants were rare, appearing only in one single allele in the KHV population. The G4500 call set included 39,485 variants (58.8%) that had not been reported in the KHV call set. Among them, 30,681 (45.7%) were found in the dbSNP database and the remaining 8,804 (13.1%) were novel. Most of the novel variants had allele frequencies less than 5% (Figure 1b).

**Figure 1.**
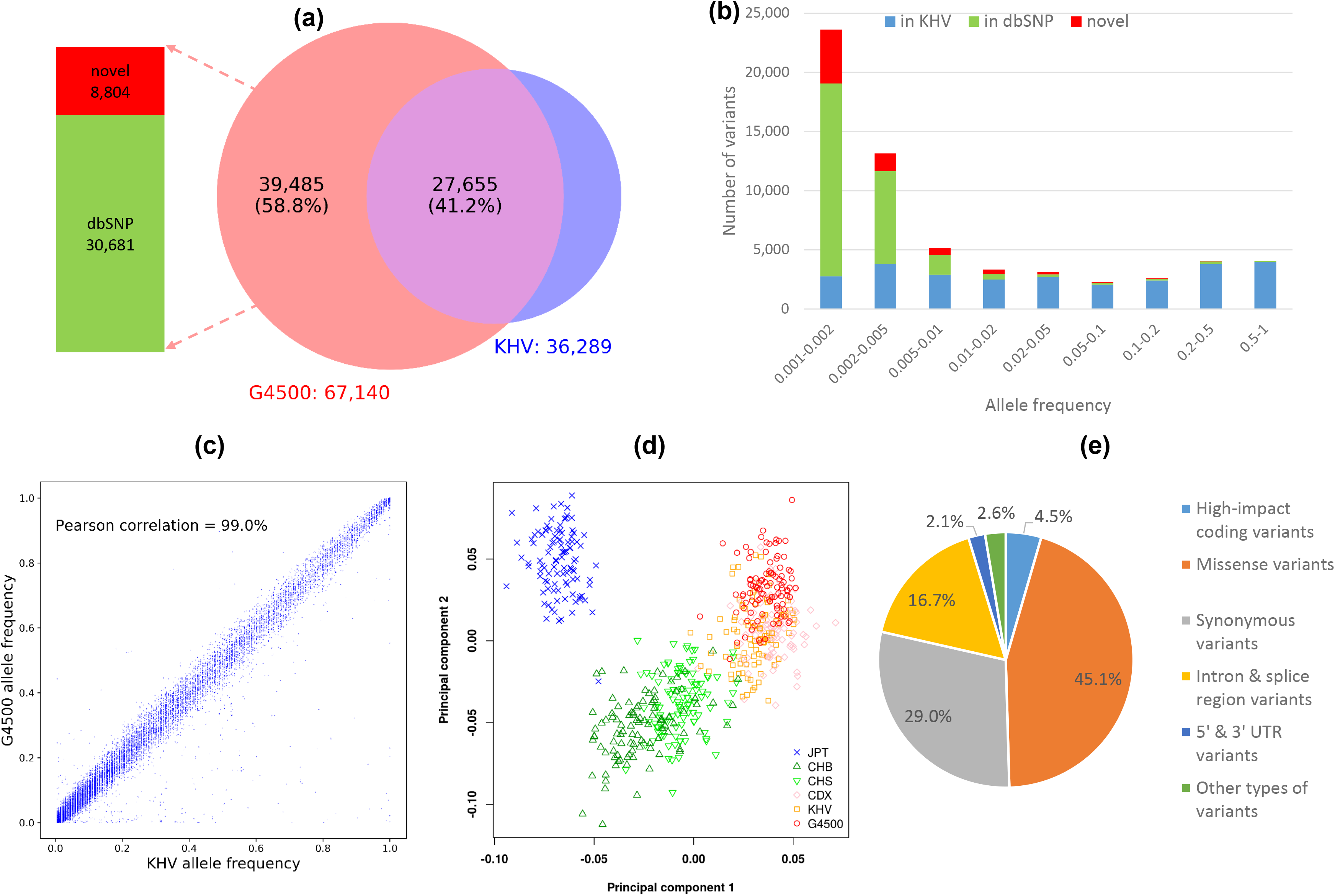
Summary of the G4500 variant call set. (a) Comparison of G4500, KHV, and dbSNP. (b) Allele frequency distribution of the G4500 call set. (c) Comparison of allele frequency between G4500 and KHV. (d) Principal component analysis of the G4500 call set and other East Asia populations (JPT: Japanese in Tokyo, Japan; CHB: Han Chinese in Beijing, China; CHS: Southern Han Chinese; CDX: Chinese Dai in Xishuangbanna, China; KHV: Kinh in Ho Chi Minh City, Vietnam). (e) Distribution of variant consequences of the G4500 call set (high-impact: stop-gained, stop-lost, start-lost, frameshift, splice receptor, and splice donor).

We also performed principal component analysis (PCA) on the G4500 population and other East Asia populations from the 1000 Genome Project (JPT: Japanese in Tokyo, Japan; CHB: Han Chinese in Beijing, China; CHS: Southern Han Chinese; CDX: Chinese Dai in Xishuangbanna, China; KHV: Kinh in Ho Chi Minh City, Vietnam). Overall, Figure 1d shows that the PCA clustering of the populations was consistent with their respective geographic locations. The G4500 and KHV populations closely clustered together as both represented the Vietnamese population. They were also located closer to the CDX population than to the CHS, CHB, and JPT populations, agreeing with the respective geographical distances.

We then used Variant Effect Predictor (VEP [18]) to predict potential effects of variants in the G4500 call set (Figure 1e). Majority of them were missense variants (45.1%), followed by synonymous variants (29.0%), and intron or splice region variants (16.7%). Notably, 4.5% of the variants were predicted to have high-impact consequences, including stop-gained, stop-lost, start-lost, frameshift, splice receptor and splice donor. Those high-impact variants may lead to protein truncation and are critical for clinical interpretation, as we shall show in the next sections.

### Carrier frequencies of genetic diseases in the Vietnamese population

We annotated the G4500 variant call set against the ClinVar database to identify pathogenic variants, genes, associated diseases, and estimated their carrier frequencies in the Vietnamese population. We found 21,151 variants with ClinVar annotations, and among them, 158 variants had been reviewed as “Pathogenic” or “Likely pathogenic”. These 158 variants were located on 116 genes: 84 genes were associated with autosomal recessive (AR) diseases, 18 genes with autosomal dominant (AD) diseases, 9 genes with both AD and AR diseases, one gene with X-linked dominant disease (XLD) and one gene with X-linked recessive disease (XLR). In this study, we focused on 114 pathogenic variants on 85 genes that were associated with recessive diseases (84 AR and one XLR).

Twenty-three individuals in our cohort were identified as homozygous or compound heterozygous carriers for 5 genes associated with recessive diseases, including *GJB2* (n=12), *HFE* (n=5), *VPS13B* (n=4), *CBS* (n=1), and *GBA* (n=1) (Supplementary Table S2). Since our cohort data was obtained from pre-existing hospital records rather than a randomized study design, we took a conservative approach and excluded these 23 individuals before calculating the carrier frequencies of the respective genes and diseases. Overall, the carrier frequencies were reduced by 0.1%-1% by this exclusion (Supplementary Table S3).

Figure 2 shows a summary of 114 pathogenic variants on 85 genes and associated recessive diseases identified from our G4500 dataset. The complete details are provided in Supplementary Table S4. As shown in Figure 2a, majority (54%) of these variants were protein-truncating (including stop gained, frameshift, splice acceptor or donor), followed by missense variants (41%). While most of the 85 genes only had one pathogenic variant, 20 of them (23.5%) had at least two pathogenic variants per gene (Figure 2b), such as *GAA* (5 variants), *GJB2* and *HBB* (3 variants each), *VPS13B* (2 variants), etc (Supplementary Table S4). By taking into account all pathogenic variants of each gene, our study provided more accurate estimates of disease carrier frequencies than a targeted genotyping approach that only focused on major variants [2]. The carrier frequency distribution is presented in Figure 2c. 17/85 genes (20%) were estimated to have carrier frequencies of more than 1% (1 in 100), among which seven diseases appeared in more than 2% (1 in 50), including three appeared in more than 5% (1 in 20) of the Vietnamese population.

**Figure 2.**
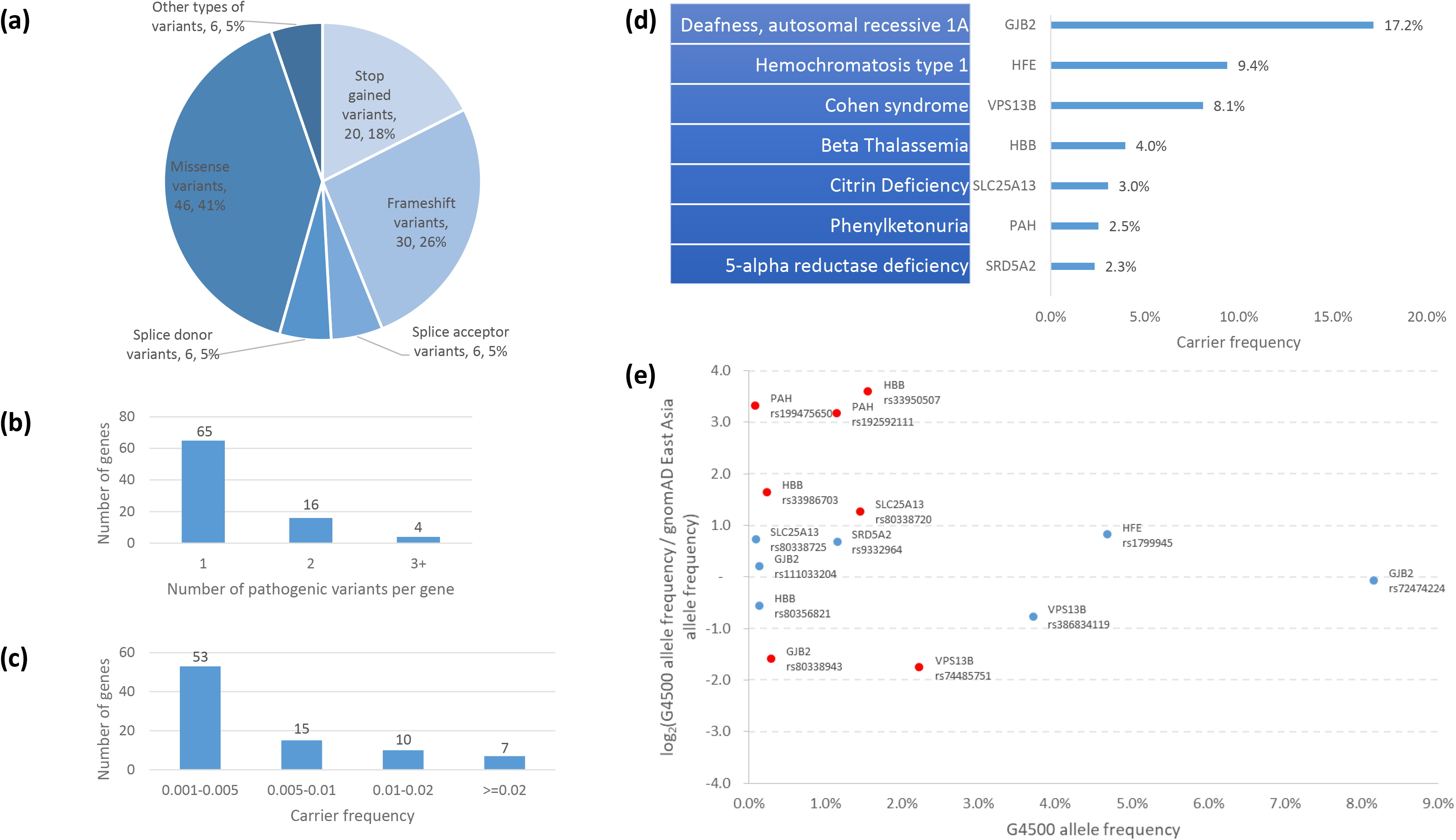
Summary of pathogenic variants, genes, and associated recessive diseases identified from the G4500 dataset. (a) Distribution of coding consequences of pathogenic variants. (b) Distribution of pathogenic variants per gene. (c) Distribution of carrier frequencies of pathogenic genes. (d) Top seven diseases-genes with carrier frequencies of more than 2%. (e) Allele frequencies of pathogenic variants of the top seven diseases-genes in the Vietnamese G4500 population (x-axis) and how they are compared to the frequencies in the East Asia population (y-axis). Some genes may have multiple variants, e.g. *GJB2* has three variants. Red points indicate variants with allele frequencies different by more than two folds between the two populations (i.e. log_2_ fold change is less than −1 or greater than 1). The details of these variants are provided in Table 2.

Figure 2d shows the top seven genes and associated recessive diseases with carrier frequencies of more than 2% (1 in 50) in the Vietnamese population. Deafness, autosomal recessive 1A associated with gene *GJB2* was the most prevalent disorder with a carrier frequency of 17.2% (1 in 6). The prevalence of *GJB2*, in particular, the SNP rs72474224 C>T, in the Vietnamese population and other East Asian populations, as compared to Western populations, had been reported previously in [9]. Two other autosomal recessive diseases were found with relatively high carrier frequencies, including hemochromatosis type 1 (*HFE*, 9.4% or 1 in 11) and Cohen syndrome (*VPS13B*, 8.1% or 1 in 12). Hemochromatosis type 1 is a metabolic disorder that causes the body to absorb too much iron (iron overload). Cohen syndrome is a multisystem disorder characterized by many clinical features, including developmental delay, intellectual disability and facial dysmorphis. Both diseases are common genetic disorders among Western populations, but we found that they appeared less frequently in the Vietnamese population (Table 2). We also observed three other disorders that are among the most commonly encountered diseases by local medical doctors in Vietnam, including Beta-thalassemia (*HBB*, 4% or 1 in 25), citrin deficiency (*SLC25A13*, 3% or 1 in 33), and phenylketonuria (*PAH*, 2.5% or 1 in 40).

**Table 2.**
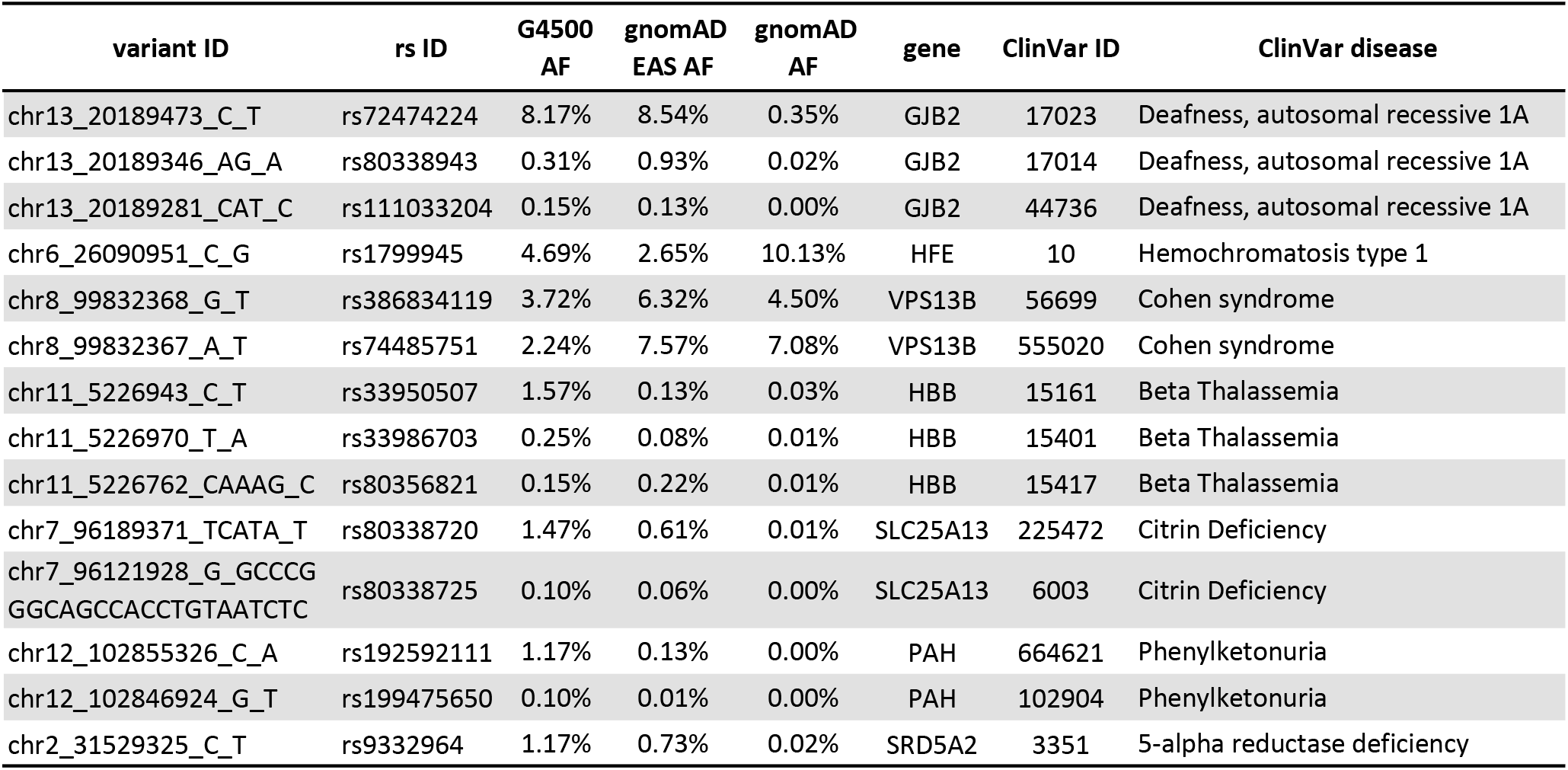
Pathogenic variants of the seven most prevalent diseases-genes in the Vietnamese population. (AF: allele frequency; CF: carrier frequency; EAS: East Asia)

### Beta-thalassemia, Citrin Deficiency, and Phenylketonuria

We further compared the allele frequencies of pathogenic variants of the top seven diseases-genes between the Vietnamese, the East Asia, and the global populations (gnomAD [20]). Figure 2e and Table 2 show that several pathogenic variants appeared 2-12 times more frequent in the Vietnamese population, especially for three diseases Beta-thalassemia, citrin deficiency and phenylketonuria. Beta-thalassemia is a blood disorder that reduces the production of hemoglobin; its major type can lead to severe or life-threatening outcomes and requires frequent blood transfusions for red blood cell supply. The prevalence and severe consequences of Beta-thalassemia is well-known among the Vietnamese population, yet no research has been done to study its genetic patterns in the population. Here we found that the allele frequency of the SNP rs33950507 C>T in gene *HBB* was 12 times higher in the Vietnamese population than in the East Asia population (1.57% and 0.13%, respectively). Furthermore, rs33950507 and two other pathogenic variants in gene *HBB* collectively contributed to a carrier frequency of 4% (1 in 25) for Beta-thalassemia in the Vietnamese population. The global carrier frequency of Beta-thalassemia had been estimated previously as 0.7% (1 in 143), i.e. 5.7 times lower than in the Vietnamese population [2].

Another two SNPs, rs192592111 C>A and rs199475650 G>T, in gene *PAH* and associated with phenylketonuria were also found to have allele frequencies 9 times higher in the Vietnamese population than in the East Asia population (Figure 2e, Table 2). Phenylketonuria is a metabolic disorder that causes phenylalanine to build up in the body, and if not treated, may lead to intellectual disability and other serious health problems. This disease is a very rare genetic condition in the world with a carrier frequency of 0.7%, mostly observed in Southern Europe or Hispanic, but not among the East Asia population [2]. However, we found two of its variants and estimated that its carrier frequency was 2.5% (1 in 40) in the Vietnamese population.

Similarly, we found two SNPs rs80338720 and rs80338725 in gene *SLC25A13* that were associated with citrin deficiency, and their respective allele frequencies were 2.4 times and 1.6 times higher in the Vietnamese population than in the East Asia population. The total carrier frequency of citrin deficiency was estimated as 3% (1 in 33) in the Vietnamese population, which was in line with recent results for South East Asian populations in Singapore [3]. Citrin deficiency is a metabolic disorder that manifests in newborns as neonatal intrahepatic cholestasis or in adulthood as recurrent hyperammonemia with neuropsychiatric symptoms in citrullinemia type II. Without appropriate treatment, severe liver problems may develop and require liver transplantation.

In addition to the top seven genes with carrier frequencies of more than 2% (1 in 50), ten other genes had carrier frequencies of at least 1% (1 in 100), and the remaining 68 genes had carrier frequencies of less than 1% in the Vietnamese population. The complete profile of pathogenic variants, genes, recessive diseases, and their frequencies in the Vietnamese population is provided in Supplementary Table S4. Some other examples of high carrier frequencies include Pompe disease (*GAA*, 1.9% or 1 in 52), Zellweger syndrome (*PEX1*, 1.6% or 1 in 62), Stargardt disease (*ABCA4*, 1.3% or 1 in 76), Krabbe disease (*GALC*, 1.3% or 1 in 76), Bestrophinopathy, autosomal recessive (*BEST1*, 1.1% or 1 in 90), and Wilson disease (*ATP7B*, 0.9% or 1 in 110).

### Identifying new pathogenic variants for the Vietnamese population

We next attempted to identify new pathogenic variants for the Vietnamese population from the G4500 call set. We focused on the variants that were predicted by VEP to have high-impact consequences but had not been reported in ClinVar. We identified 131 variants that may cause protein truncation, including stop-gained, stop-lost, start-lost, frameshift, and splice receptor or donor disruptions. Their distribution is presented in Figure 3a. We then manually reviewed these variants according to the American College of Medical Genetics (ACMG) classification guidelines [21] and classified seven of them as “Pathogenic” and three as “Likely pathogenic” variants. Their details are presented in Figure 3b and Supplementary Table S5.

**Figure 3.**
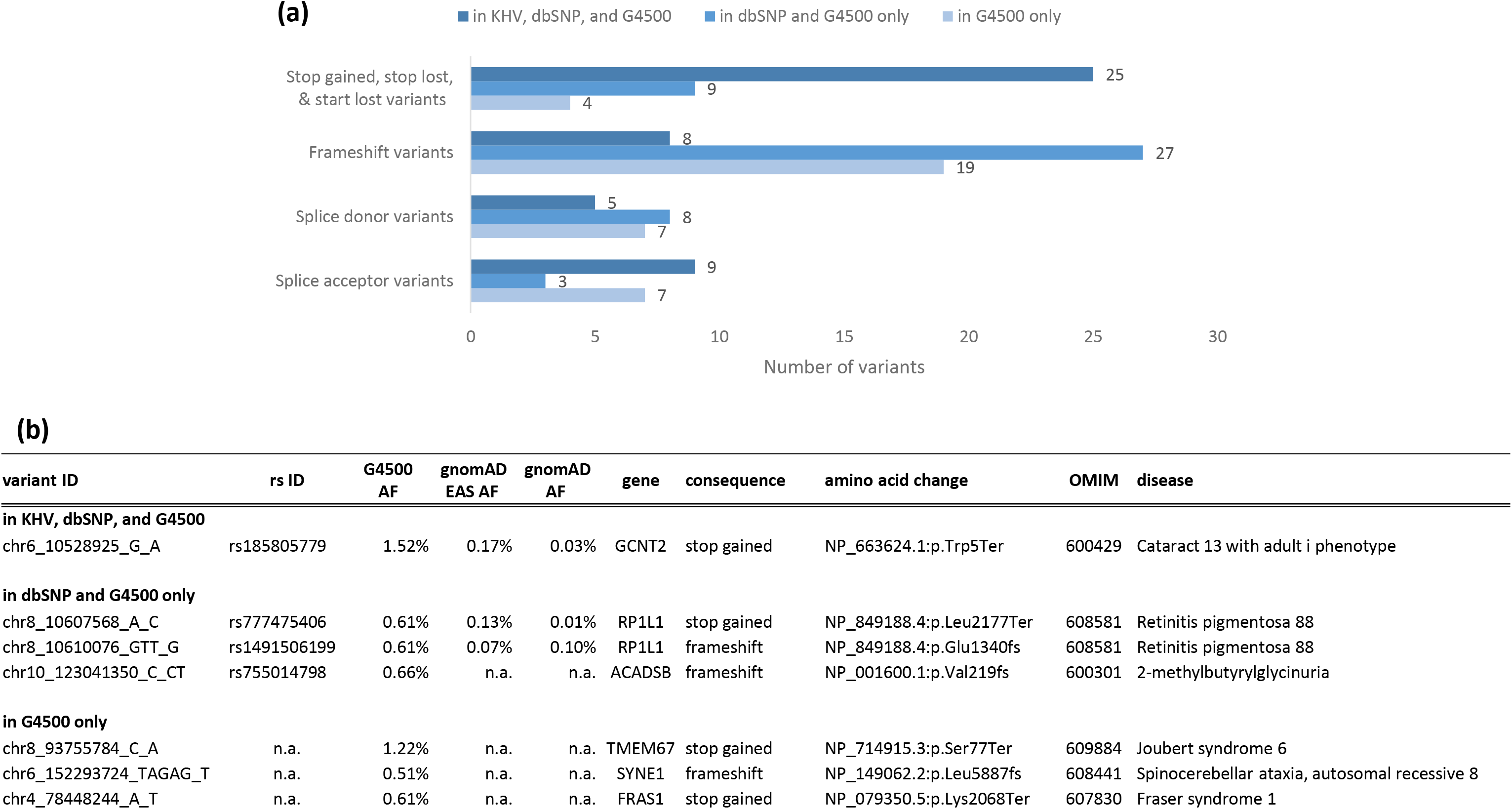
(a) Distribution of variants identified from the G4500 dataset that had high-impact consequences but had not been reported in the ClinVar database. (b) Seven new pathogenic variants that we selected from (a), reviewed, and classified as “pathogenic” according to the ACMG guidelines. (AF: allele frequency; EAS: East Asia; ACMG: American College of Medical Genetics).

The seven new pathogenic variants include four stop-gained and three frameshift variants that are rare or not present in public databases. In particular, four of them were found in gnomAD with global allele frequencies ≤0.1% and three of them were only found in our G4500 dataset. Their allele frequencies in the Vietnamese population were several times higher than in the East Asia and the world populations. For instance, the SNP rs185805779 G>A had allele frequencies of 1.52%, 0.17%, and 0.03% in the Vietnamese, the East Asia, and the global populations, respectively (Figure 3b). This stop-gained variant in gene *GCNT2* leads to a premature termination codon p.Trp5Ter at the beginning of the protein NP_663624.1 and disrupts this whole protein. Similar nonsense, loss-of-function variants in gene *GCNT2* had been reported as pathogenic and associated with the cataract 13 with adult i phenotype, an autosomal recessive disorder of i and I antigens in blood that may lead to congenital cataract (OMIM 600429). Thus, we classified this variant as pathogenic (evidence categories PVS1, PM2 and PM4 in ACMG guidelines).

Notably, we identified three novel pathogenic variants that had never been reported before in any databases. In particular, the SNP chr8:93755784 C>A in gene *TMEM67* is a stop-gained variant that causes a premature termination codon p.Ser77Ter on the protein NP_714915.3. Two other stop-gained, loss-of-function variants on this gene and its protein had been reported in ClinVar as pathogenic, including ClinVar 506012 (NP_714915.3:p.Arg172Ter) and ClinVar 1376 (NP_714915.3:p.Arg208Ter). Note that the mutated amino acid of the new SNP is located at position 77 and hence results in a shorter truncated protein than the other two mutations, causing even more severe damages. We classified this new SNP as pathogenic for *TMEM67-* associated Joubert syndrome (OMIM 609884).

Another novel stop-gained variant that we classified as pathogenic was the SNP chr4:78448244 A>T in gene *FRAS1*, which causes a premature termination codon p.Lys2068Ter on the protein NP_079350.5. Note that a missense variant, rs1578330963 A>G, had been reported at the same location in dbSNP for the Korean population [22]. Two other stop-gained, loss-of-function variants in *FRAS1* and NP_079350.5 had been reported in ClinVar as pathogenic, including ClinVar 197861 (NP_079350.5:p.Arg124Ter) and ClinVar 435260 (NP_079350.5:p.Gln907Ter). Thus, we classified the new SNP as pathogenic for *FRAS1*-associated Fraser syndrome 1 (OMIM 607830). Fraser syndrome is a rare genetic disorder characterized by cryptophthalmos, cutaneous syndactyly, and abnormalities of the genitalia and the urinary tract.

Similarly, we classified a novel deletion variant, chr6:152293724 TAGAG>T, as pathogenic for *SYNE1*-associated Spinocerebellar ataxia-8. This variant causes a frameshift p.Leu5887fs on the protein NP_149062.2, for which several loss-of-function frameshift variants had been reported as pathogenic (ClinVar IDs 204299, 436905, 199228). Spinocerebellar ataxia-8 is a slowly progressive neurodegenerative disorder characterized by gait ataxia and other cerebellar signs, such as nystagmus and dysarthria (OMIM 608441).

Last but not least, we classified three new splice acceptor or donor variants as likely pathogenic (Supplementary Table S5). These variants were predicted to disrupt mRNA splicing and result in an absent or disrupted protein product. They were not found or appeared at less than 0.01% frequency in gnomAD. Similar splice acceptor or donor variants on the same genes had been reported as pathogenic or likely pathogenic. For instance, the SNP rs1183832067 A>C is a splice donor in gene *RFX5*, and we found that its corresponding splice acceptor rs748270285 G>A for the same exon 6 of transcript NM_001025603.2 had been reported as pathogenic for Bare lymphocyte syndrome, type II, complementation group c (ClinVar 7646). Since more data is needed to establish the pathogenicity, we classified the three splice acceptor or donor variants in Supplementary Table S5 as likely pathogenic (evidence categories PVS1 and PM2 in ACMG guidelines).

## DISCUSSION

In this paper, we analyzed the clinical exome sequencing data of 4,503 genes obtained from a cohort of 985 individuals to study recessive diseases in the Vietnamese population. We identified a comprehensive variant call set named G4500 that includes 61,327 SNPs and 5,813 indels. We showed that the G4500 variant call set accurately represented the genetic characteristics of the Vietnamese population and also demonstrated how they are related to other East Asia populations.

Most importantly, our work is the first study that provided a comprehensive picture of 85 most common recessive diseases and their prevalence in the Vietnamese population. Among them, seven diseases had carrier frequencies of more than 2% (1 in 50) and ten diseases had carrier frequencies of at least 1% (1 in 100). For each disease, we provided complete details of its pathogenic variants, gene, and carrier frequency in the Vietnamese population as compared to other populations. For instance, *GJB2*-associated deafness autosomal recessive was the most prevalent disorder with a carrier frequency of 17.2% and consisted of three pathogenic variants. Notably, we found three diseases that were specific to the Vietnamese population with carrier frequencies of several times higher than in other East Asia or the world populations, including Beta-thalassemia (*HBB*, 4% or 1 in 25), citrin deficiency (*SLC25A13*, 3% or 1 in 33), and phenylketonuria (PAH, 2.5% or 1 in 40).

We also discovered seven new pathogenic and three new likely pathogenic variants that had not been reported in ClinVar. These new variants were associated with nine autosomal recessive diseases in autoimmune, hematology, ophthalmology, and neurology. Notably, two new pathogenic variants revealed much higher carrier frequencies of *TMEM67*-associated Joubert syndrome and *GCNT2*-associated cataract 13 with adult i phenotype in the Vietnamese population (2.64% and 3.04%, respectively) than previously estimated. Some of these variants and diseases might also appear in other populations at extremely low frequencies, e.g. *GCNT2*-associated cataract 13 with adult i phenotype and *RP1L1*-associated retinitis pigmentosa, thus representing a great resource for further studies. We also discussed how these new variants were related to previously reported pathogenic variants on the corresponding genes and proteins.

One limitation of this study was that our cohort was sampled from pre-existing hospital records rather than a randomized study design. To remove potential bias in our estimation of allele and carrier frequencies due to this type of sampling, we took a conservative approach by considering only recessive diseases and excluding 23 individuals identified as homozygous or compound heterozygous carriers for 5 genes. Thus, our estimated carrier frequencies for these 5 genes and diseases may be considered as lower bounds. A more properly designed study with sufficiently large dataset could offer a more accurate representative of the Vietnamese population.

In conclusion, our study has significantly improved the knowledgebase and the practice of medical genetics in Vietnam in many aspects. Our findings offer a great resource to inform local public health policies to understand and better align with the specific landscape of genetic diseases in the Vietnamese population. Carrier or newborn genetic screening programs can be re-designed for cost-effectiveness and comprehensiveness. The results also help clarify and expand existing knowledge of popular inherited diseases in the local population by providing the extra dimension of molecular genetic information. By demonstrating the underlying fundamental role of genetics in inherited diseases, our work also contributes to the development of genetics education, genetics counseling, and genetics screening among the local population. The identification of three inherited diseases specific to the Vietnamese population affirms the necessity of population-specific genetic studies and that larger and more comprehensive population genetic studies dedicated to the Vietnamese population are highly desired.

## Ethics approval and consent to participate

The study was approved by the institutional ethics committee of the University of Medicine and Pharmacy, Ho Chi Minh city, Vietnam. The study has followed the guidelines set by the University of Medicine and Pharmacy, Ho Chi Minh city, Vietnam, in handling human genetic data of the participants. The participants have approved and given written informed consent to the anonymous re-use of their genomic data for this study.

## Consent for publication

All authors have read and approved the manuscript for publication.

## Availability of data and materials

The G4500 variant call set is available upon reasonable request to the corresponding authors, subject to our policy of data privacy.

## Authors’ contributions

THNT, HST, LPH, THLN, NTT, THNT, VTN, BHHN, NMP. KHTN, HDLN, MTTQ, TPTN, DVT, QTTN recruited patients and performed clinical analysis.

HNN, TTTD, NVL, VUT, PCTN, DKT, HTN, LPD, designed experiments and analyzed data.

NHT, HG, MDP designed the experiments, analyzed the data and wrote the manuscript.

HNN supervised the project.

